# An atomic-scale view at the composition of amyloid-beta fibrils by atom probe tomography

**DOI:** 10.1101/351973

**Authors:** Kristiane A.K. Rusitzka, Leigh T. Stephenson, Agnieszka Szczepaniak, Lothar Gremer, Dierk Raabe, Dieter Willbold, Baptiste Gault

## Abstract

Amyloid-beta (A*β*) proteins play an important role in a number of neurodegenerative diseases. A*β* is found in senile plaques in brains of Alzeimer’s disease patients. The 42 residues of the monomer form dimers which stack to fibrils gaining several micrometers in length. Using A*β* fibrils with ^13^C and ^15^N marker substitution, we developed an innovative approach to obtain insights to structural and chemical information of the protein. We deposited the modified protein fibrils to pre-sharped aluminium needles with >100-nm apex diameters and, using the position-sensitive mass-to-charge spectrometry technique of atom probe tomography, we acquired the chemically-resolved three dimensional information for every detected ion evaporated in small fragments from the protein. We also discuss the influence of experimental parameters such as pulse energy and pulse frequency of the used Laser beam which lead to differences in the size of the gained fragments, developing the capability of localising metal atom within A*β* plaques.

## Introduction

Amyloid-beta (A*β*) proteins are involved in a number of neurodegenerative diseases^1–5^. Notably, it plays an important role in the progress of Alzheimer’s disease^6,7^. A*β* peptides are formed by sequential cleavage of the amyloid precursor protein (APP) by *β*-secretase and the γ-secretase complex^8^. In the progress of Alzheimer’s diseases, these proteins can fibrillate and assemble forming extracellular plaques in inner organs and especially the brain^9–12^. A*β* occurs in different amino acid sequence-length and the plaques found in brains of Alzheimer-affected patients are most dominantly formed by A*β*1-42^13–16^. Over several years, various studies have repeatedly indicated that various metals ions may be associated with Alzheimer’s disease, e.g. copper, iron and zinc^17,18^. The different metal ions may lead to a change in the conformation of the A*β* protein^18,19^.

For understanding the biological function of proteins and to aid designing medical drugs, it is important to know the protein structure as well as possible and to acquire chemistry of the structural components. Therefore, a mixture of different techniques is usually used to complement each other, e.g. cryo-electron microscopy (Cryo-EM) density maps are often supplemented with nuclear magnetic resonance (NMR) or crystallography data.

Depending on the protocol used for in vitro studies, a wide range of different morphologies of the fibrils can be found. For this reason, a large number of different structure models has been published over the last years^20^, which have been discovered using different methods such as nuclear magnetic resonance (NMR)^21–27^, atomic force microscopy (AFM)^28^, cryo-EM^29–33^, scanning transmission electron microscopy (STEM)^34^, x-ray crystallography^35–37^or a combination of those methods. Different states of dimerization as well as structural orientation of the proteins with diameters in the range of 7–14 nm were examined.

A possible complementary technique is atom probe tomography (APT). APT which is well established in the materials sciences is the chemically most sensitive and highest resolving probing technique used to characterize microstructures and local compositions of complex metallic alloys and semiconductors down to near atomic scale^38,39^. An APT specimen is fashioned into an approximately 100-nm radius needle and is biased to a high standing voltage with additional high-voltage or laser pulsing that triggers controlled field evaporation, a process whereby surface atoms are ionized in elemental or molecular form. The ions are then projected towards a time-resolved, single-particle detector comprising a stack of multichannel plates paired with a delay-line detector. The ion’s time-of-flight and detector impact position provides the raw data that can be processed to build a three-dimensional reconstruction by means of a reverse-projection protocol. APT offers sub-nanometer resolution and the elemental identity of every individual ion^39–44^. APT has the potential to assist in understanding chemical gradients or unambiguously identify the presence of metals atoms and clusters, which could significantly advance understanding protein structure and function. APT has also proven successful when applied to biological materials. Previous studies have demonstrated near-atomic resolution of chemically–resolved data of bulk bio-mineralized hard biological matter such as teeth and bone^45–51^. Progress was also made on more soft materials such as cells^52,53^and also with ferritin as example for proteins^54–56^. In early designs of the atom probe microscope, single amino acids, nucleic acids and other polymers were successfully analyzed^57–60^.

Due to the still unsolved question of the presence of metal ions in naturally occurring A*β* fibrils, APT provides an opportunity to obtain structural information as well as the precise chemical identity of the ions. This different approach can also open the possibility to examine several other proteins which are associated to metal ions. Here, we used the same fibrils that were also the basis of a recently published 3D-reconstruction, made using data from cryo-EM and solid-state NMR^33^. The structure described therein refers to fibrils with a diameter of 7 nm. Instead of natural isotopes, C_13_ and N_15_ were incorporated in the protein. We present preliminary exploration of the possible application of APT to investigate the details of the elemental distribution within A*β* fibrils and discuss the influence of experimental parameters such as pulse energy and UV-laser pulse frequency on the data quality.

The aim of our work is to find a protocol to analyze A*β* fibrils by APT and know the influence of the main experimental parameters so as to maximize the accuracy of the measurements. We establish a step-by-step protocol with a list of parameters that need to be optimized, so that we can move ahead and start a systematic investigation, bearing in mind that the future target will be on detecting small quantities of metal ions. In this study we investigated the success of sample preparation based on the incubation time of the protein to the pre-sharped specimen as well as the influence of pulse energy of the laser in a range of 10 pJ to 40 pJ and pulse frequency of the laser in a range of 125 kHz to 500 kHz.

## Results and Discussion

### Specimen Preparation

The first aspect we address is the preparation of specimens appropriate for reproducible APT analysis of amyloid-beta fibrils. To avoid damaging the proteins during preparation, as perhaps from plasma FIB milling, we deposited the protein on a pre-sharpened Al-specimen (Fig 1A). Gremer et al. have shown that the stability of A*β* is not affected by air drying the protein^33^. By air drying the sample, issues like brittleness through dehydrating agents or fixatives could be avoided. With a diameter of 7 nm and several micrometers in length^33^, it was not possible to control the results of deposition without staining in SEM or TEM.

### Optimization of sample preparation

A schematic overview of the sample preparation is given in Fig 1. Ions, evaporated from the specimen impact coordinate-sensitive detector (Fig 1B), while a mass-to-charge spectrum is created (Fig 1C) based on their time-of-flight (Fig 1C). In all APT experiments, the measurements exhibited enough chemical contrast to distinguish the organic parts and the supporting Al-specimen (Fig 1D). The thickness of the protein layer on the Al-needle should increase with longer incubation times. However, the primary target was to investigate individual fibrils at highest possible chemical precision and hence a thick layer was not desirable. Besides, proteins are relatively poor electrical conductors^52^. Therefore, a thicker protein layer will likely require a higher applied voltage to induce the surface field necessary to produce and evaporate ions thus possibly increasing the risk of specimen fracture. Most specimens started emission between 2-3 kV and from these we acquired between one and four million ions per APT tip probed. Longer incubation times demonstrated a higher likelihood of specimen fracture early in an APT measurement. Several specimens, especially those with longer deposition times of the protein, fractured without any even emitting ions at a high voltage.

### APT measurement

To provide a smooth and Xe-free surface, we did a short APT measurement (1-2 mio ions) before deposition of the protein. In those specimen which were plasma FIB finish milled at energies above 16 kV, 90 pA, Xe was was implanted and observed in the mass-to-charge spectra (Fig 2).

Different from metals and semiconductors, biological materials are characterized by a complex bonding energy landscape, ranging from very weak Van Der Waals bondings (0.1 eV) over ionic to several eV strong covalent bonds. Their integrity and decomposition when exposed to high voltage fields and laser induced temperature increase can be very sensitive to the APT experiment conditions and so we were cautious when setting the laser pulse energy to minimize thermal damage, this being important to achieve well-behaved field evaporation^45,52,54,55,57^. We varied the laser pulse energy from 10 pJ up to 40 pJ (Fig 3) and the repetition rate between 125 kHz to 500 kHz (Fig 4) to explore their respective impact upon the quality of the mass-to-charge spectrum, particularly with respect to thermal tails coming from evaporation after the pulse has ended because of remaining thermal energy in the specimen and DC evaporation (described below).

**Figure 1.**
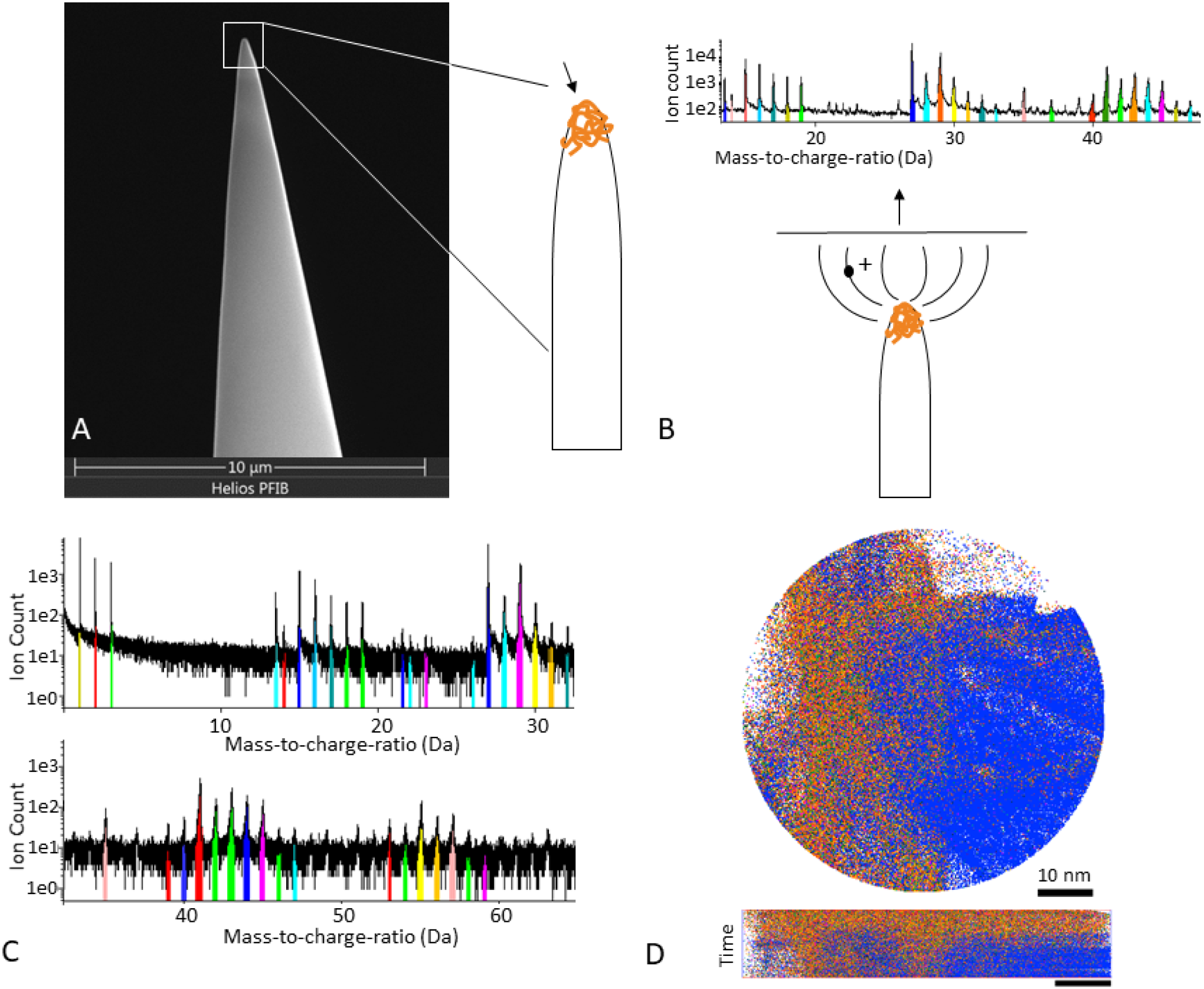
Sample preparation. A: SEM image of electropolished Al-specimen with schematic how A*β* might be deposited on Al-specimen. B: Schematic image of Al-specimen with protein. In APT, ions evaporate from the sample surface and hit a detector. Based on the time-of-flight it is possible to identify the nature of the ion and create a mass-to-charge spectrum. C: mass-to-charge spectrum from an APT measurement with ranged regions of interest which stand for different ions we examine. D: Detector map in APT measurement showing all ranged atoms in top- and side view; dark blue is 27 Da (Al), orange is 29 Da (^13^C^15^NH/^13^C_2_H_3_/^13^CO).

In mass-to-charge correlation histograms, diagonal trails with positive gradient indicated one of two mass-to-charge spectrum features in a laser-pulsed APT experiment^61^. One trail type is termed a heat tail when using laser-pulsed atom probe. This type of effect can often appear following some peaks in a one-dimensional mass-to-charge spectrum and, in a two-dimensional correlation histogram, they manifest as a smear of points from a coordinate pair (m_1_,m_2_) to greater masses. They are induced by a prolonged heating of the tip and, in the current results, they preferably appear at a laser pulse energy of 40 pJ and are seen to decrease with lowering this energy (Fig 3). At low pulse energies (10/15 pJ), only slight heat tails appear and we can accurately range many peaks in the one-dimensional mass-to-charge spectrum (Fig 3). Ranging is more difficult at the higher pulse energies (20/40 pJ) as smaller peaks can disappear, their signal vanishing in the heat tails of former peaks as well as being subject to such heat-tail spread themselves.

The second trail type is from evaporation uncorrelated with the timed laser pulse and contributes evenly to the time-of-flight signal, increasing the one-dimensional mass-to-charge spectrum background. This trail results from field evaporation events that are correlated with each other but are not time-correlated with the laser pulse, being stimulated merely from a sufficiently high specimen voltage (hence “DC” evaporation). In a two-dimensional mass-to-charge correlation plot with *m*_2_ ≥ *m*_1_, they appear as a curved lines crossing the m_2_-axis at m_1_ = 0, with the actual ion identities associated with mass-to-charge pairs (m_1_’,m_2_’), coordinates through which such curves pass. In such two-dimensional correlation plots, these DC evaporation trails can superimpose with heat tails, the latter phenomena being temporally similar. DC evaporation was evident in all experiments with pulse energies below 20 pJ and this contributed to the background in the one-dimensional mass-to-charge spectra (Fig 5). At constant detection rates, lowering the pulse energy results in an increase in specimen voltage, which increases the field experienced by surface atoms. This was evident in the change of charge-states detected; with lower pulse energies (10/ 15 pJ) higher levels of C^2+^ (6/6.5 Da) and N^2+^ (7/7.5 Da) were observed because of a higher surface field. Restoration of the peak pattern by alternation of the pulse energies was always possible and, between experiments, peaks consistently appear at the same mass-to-charge-ratio (Fig 3). A higher field also leads to smaller peaks in the detection. Beside the peak at 27 Da pertaining to Al, the most prominent peak at 10 pJ was at 29 Da (C_2_H_3_, CNH, CO), while at 20 pJ it was at 44 Da (C_3_H_5_/ CNO) and 45 Da (C_3_H_6_, CNHO, CO_2_).

**Figure 2.**
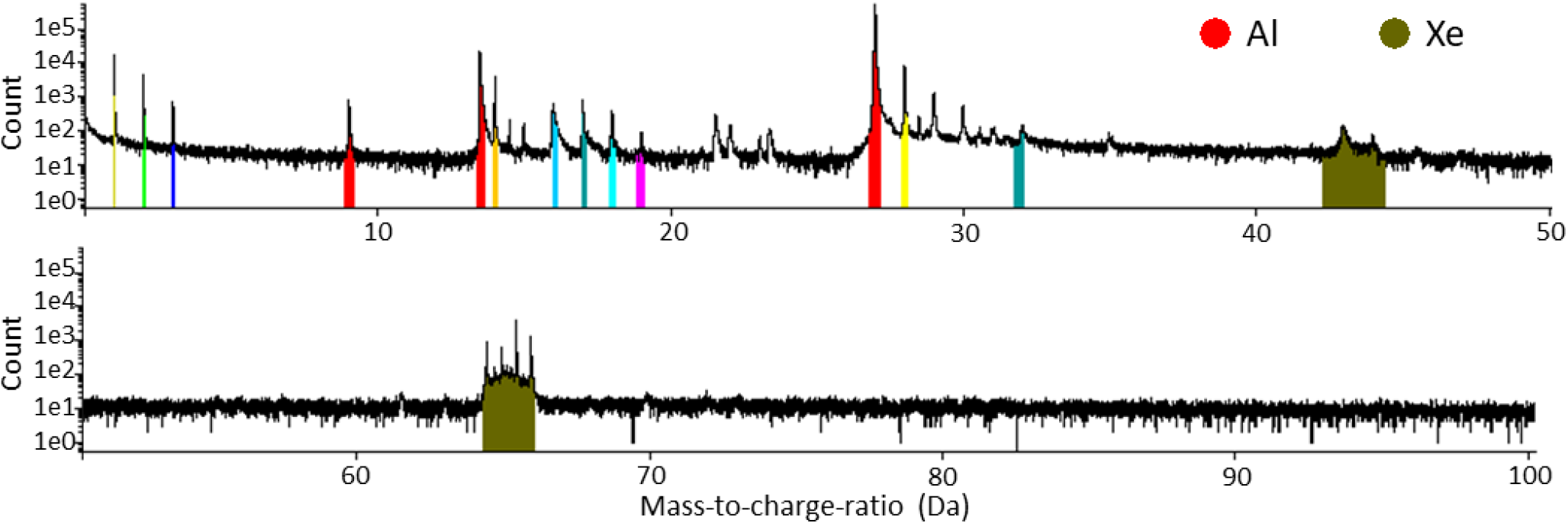
Mass-to-charge Spectra of Al-specimen with Xenon. APT measurements of Al-specimen which were sharpened with final milling parameters by PFIB higher than 16 kV and 90 pA with implemented Xe.

**Figure 3.**
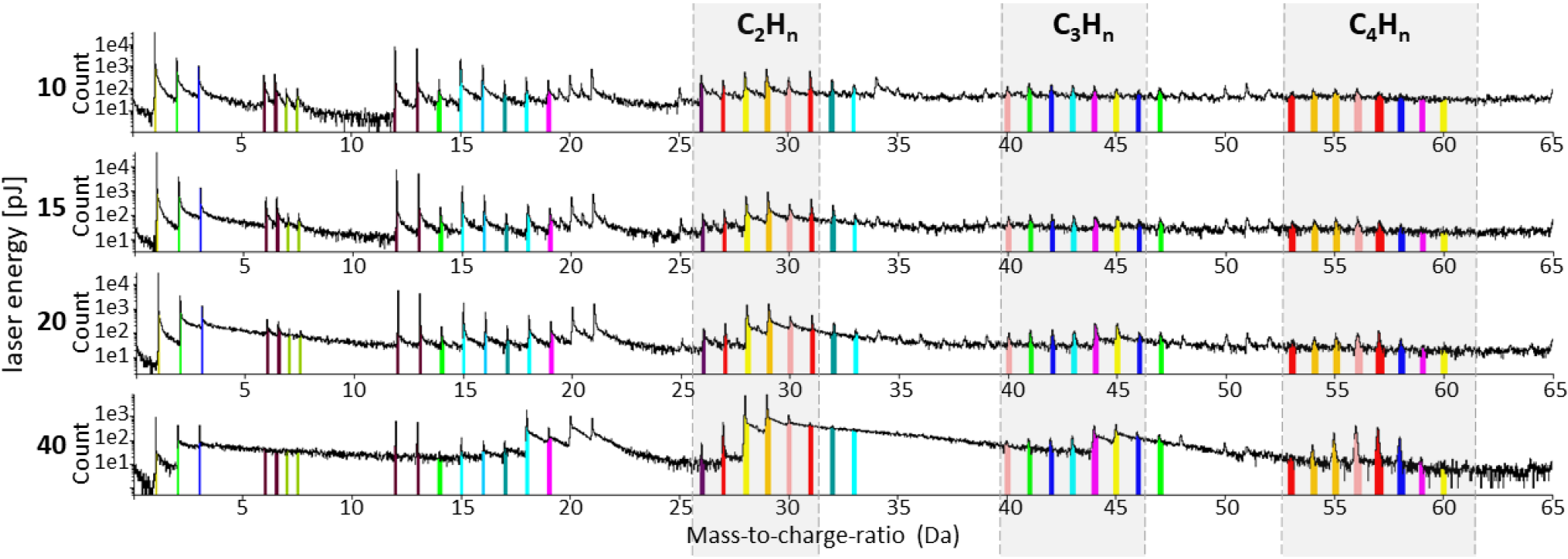
Influence of different pulse energies on mass-to-charge-ratio. Atom probe measurement with all experimental parameters constant (50 K, 125 kHz) but varying pulse energy. Four different pulse energies were tested: 10 pJ, 15, pJ, 20 pJ and 40 pJ. At 10 pJ/15 pJ peaks are good to identify. Each color band mark a different ion type composed of the protein fragments. The higher the pulse energy is, the more small peaks vanish in a smear of heat tails from the former peaks. By lowering the pulse energy again the smear vanishes and the former peak pattern appears again.

In all APT measurements, longitudinal structures of organic ions are observed. We observed that the detected spatial structure of the protein seems to prevail even at higher pulse energies (Fig 6). No noticeable changes in the structure are observed when alternating this parameter, which would perhaps simply lead to a different pattern in breaking of the amino acid bonds while field-evaporating. Changing the pulse rate and keeping the pulse energy constant at 10 pJ show slight changes in the resolution of the peaks (Fig 4).

**Figure 4.**
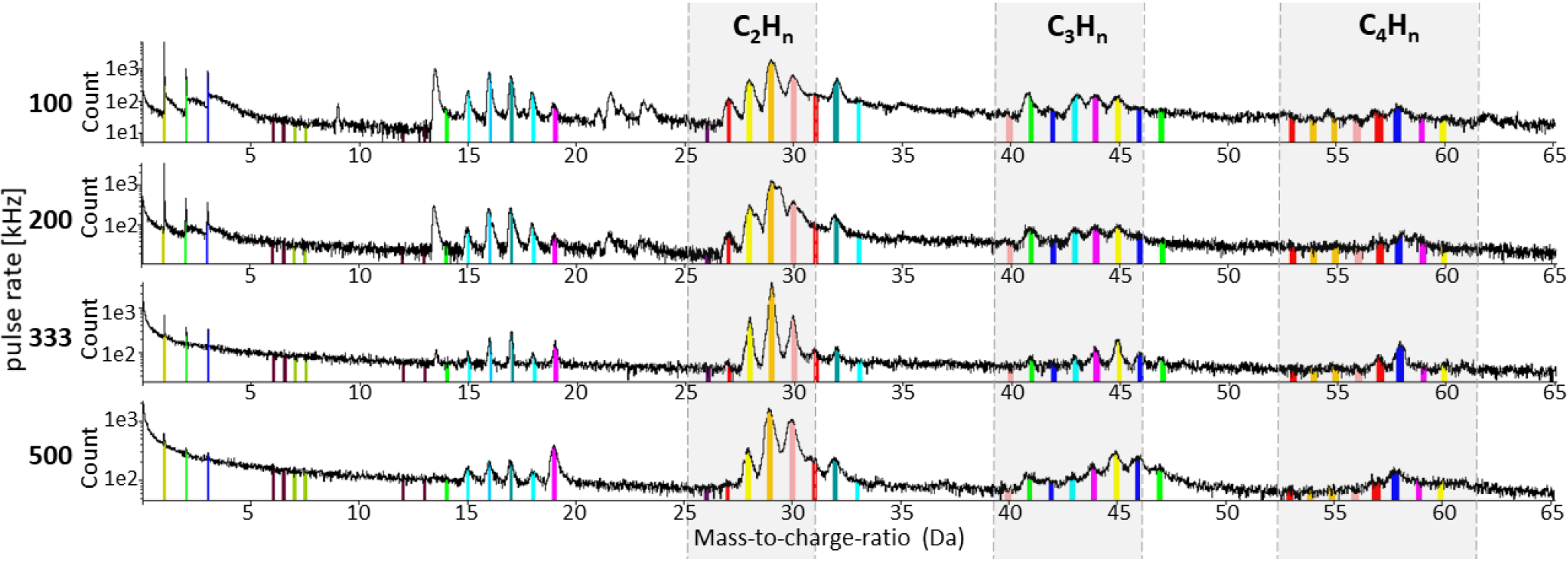
Influence of different pulse rates on mass-to-charge-ratio. Changing the pulse rate has only slight influence to the appearance of the individual peaks. Four different pulse rates were tested (100 kHz, 200 kHz, 333 kHz and 500 kHz) using a 10-pJ pulse energy. The identified ions and complex ions are ranged in different colors.

### APT mass spectrum analysis

Proteins consist of hydrogen, carbon, nitrogen, oxygen and sulfur (sulfur is part of the amino acids cysteine and methionine). The molecular formula of an amyloid-beta 1-42 monomer is C_203_H_311_N_55_O_60_S. The protein forms dimers which are stacked together to form the fibrillar structure. We expect distinct peaks at 13.00 Da (^13^C^+^), 15.00 Da (^15^N^+^) and 16.00 Da (O^+^). After most of the protein was evaporated, peaks at 9, 13.5 and 27 Da occurs which correspond to three charge states of Al. Peaks at 13 Da (^13^C^+^), 14 Da (^13^CH or ^12^CH_2_/^14^N from the solvent), 15 Da (^15^N/^13^CH_2_ or ^12^CH_3_ from the solvent), 16 Da (^14^NH_2_/O/^13^CH_3_), 17 Da (OH/^15^NH_2_/^14^NH_3_), 18 Da (H_2_O/^15^NH_3_) and 19 Da (OH^3+^) are found in all measurements. Unless specified otherwise, C and N correspond to the isotopes ^13^C and ^15^N respectively.

The peaks at 14 Da, 17 Da and 18 Da do not correspond to the longitudinal structures given by ions like CNH/C_2_H_3_/CO (29 Da). The main peaks corresponding to the organic phase are at 29-31 Da (C_2_H_n_) and 41-45 Da (C_3_H_n_). The whole data sets reveals many overlapping mass-to-charge peaks, so it is not possible to certainly identify the chemical identity of the peaks. Most of the peaks can contain more complicated combinations of elements. It is likely that the peaks between 26 Da and 32 Da correspond not only to some C_2_H_n_ but also to several C, N and H ion combinations which occur in the protein backbone. The peaks between 40 Da and 46 Da contain both C_3_H_n_, C_2_NH_n_ and CNO/CNOH. Peak overlaps like these makes it challenging to give a clear picture of the detected ion species (Tab 1).

### Image stack used as 3D-reconstruction

Based on the mass-to-charge deconvolution and fragment association outlined above we we were able to distinguish between organics which occur as different combinations of C, H, O and N and the support specimen (Al) (Fig 6). The organic compounds are mainly observed as longitudinal structures which occurred in all measurements. This would fit to our expectation as the proteins are deposited as a thin layer on the surface of the Al-specimen. The organic compounds occurred mainly in the beginning of the measurement while Al is only evaporated later in an experiment (Fig 6). Interestingly different types of ions such as CNH (29 Da) and Al (27 Da) have been detected only in specific regions, i.e. ions belonging to the protein follow the longitudinal structure, Al-ions are detected around or underneath it and some ions are randomly distributed (Fig 7), possibly due to trajectory aberrations in the evaporation and projection of ions. A*β* contains one methionine, but it was not possible to locate the single sulfur ion in it owing to the many peak overlaps and due to insufficient knowledge on the bond breaking. Also, it remains unclear at his stage if sulfur is combined with other ions while evaporating. (Tab 1).

**Figure 5.**
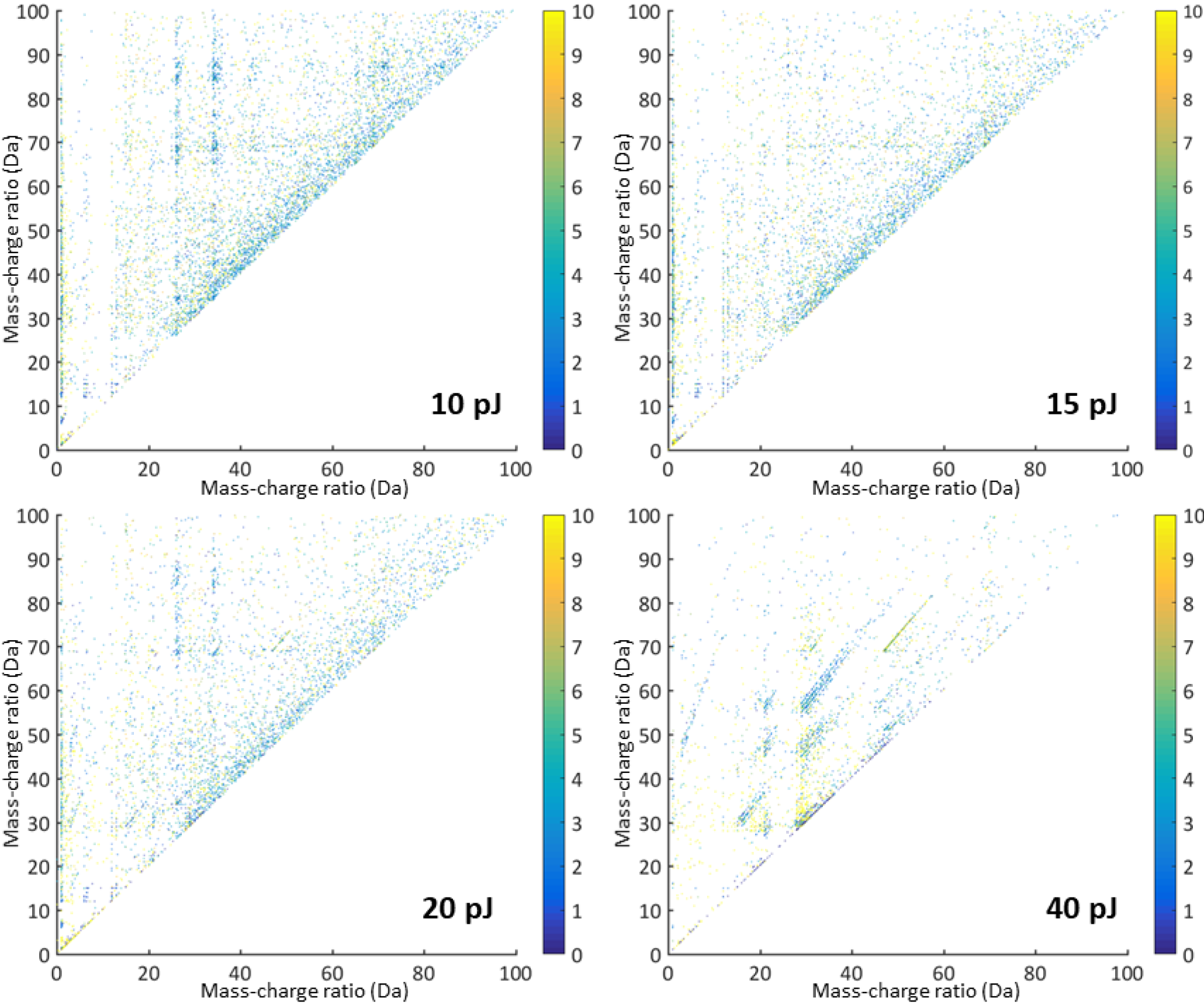
Correlation histograms. Correlation between the mass-to-charge ratios of double ion detections shown for different laser pulse energies; for the corresponding mass-to-charge spectra see Fig 3. Colormap showing the how many ions hit the detector at the same time.

## Conclusions

We studied the chemical composition of proteins that are related to Alzheimer’s disease by atom probe tomography. Firstly, we showed that A*β* fibrils can be readily deposited as a thin layer onto an Al substrate. Secondly, we demonstrated that this protein specimen could be successfully analyzed using laser-pulsed atom probe tomography. Instead of using any standard three-dimensional reconstruction protocol, we directly reconstructed image stacks from atom probe detector information to study the field evaporation sequence. This approach gave a good representation of the ion distributions and sample morphologies but, in general, in all datasets we observed an organic layer on top of the aluminum. Variation of experimental parameters (especially laser energy) allowed the acquisition of high quality mass-to-charge spectra evincing small peaks and few heat tails. For these acquisitions, 10/15-pJ laser energies were used as with higher laser energies the formation of heat tails caused the less significant mass-to-charge signals to be entirely lost. Otherwise, all mass-to-charge spectra consistently demonstrated the same prominent peaks. A definite peak assignation was difficult as the different combinations of C, H, O and N chains possible from the protein allows many isobaric peak overlaps.

Further work will consider the optimization of the parameters concerning specimen fabrication. Data mining techniques like hyperspectral analyses could also provide protein structure information. Different metals for the substrate will be trialled so as give better contrast with to ion’s evaporated from the deposited protein. In the next steps we try to identify metal ions coordinated by A*β* fibrils.

**Figure 6.**
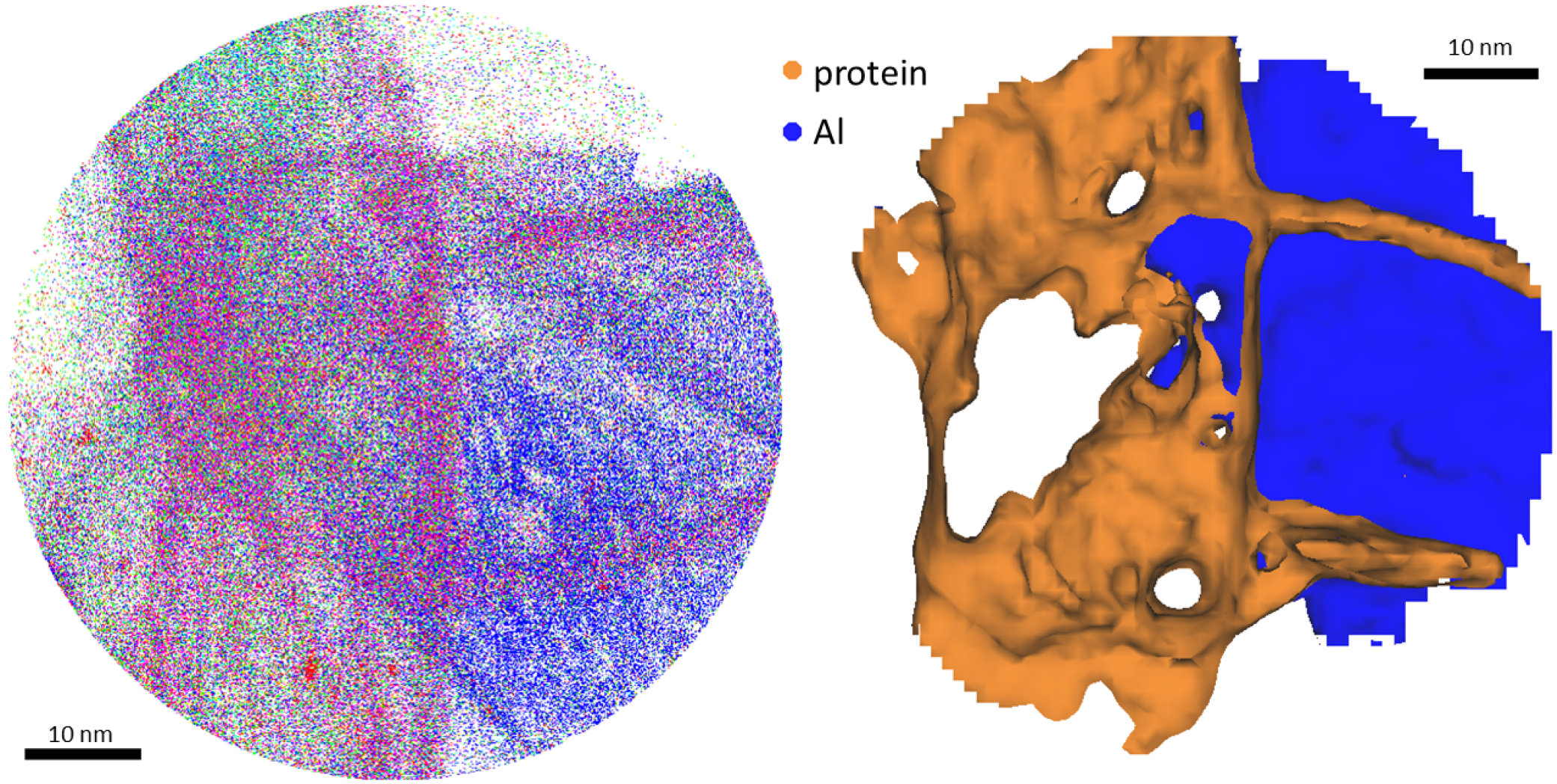
Difference between organic and inorganic parts of the specimen. Detector map showing evaporation of organic and inorganic material. A: Distribution of ions coming from the prominent peak in the mass-spectrum at 29 Da (C_2_H_3_, CNH, CO) in pink; Al underneath the protein layer marked in dark blue. Other colors are various types of ions from different fragments of the protein. B: Based on iso-surface concentration, dark blue: Aluminum from the substrate, orange: protein material.

## Methods

### Protein

Amyloid-beta 1-42 (A*β*1-42) fibrils^33^ were provided by the research group of Dieter Willbold (Heinrich Heine University Düsseldorf, Germany; Forschungszentrum Jülich, Germany). They were recombinantly expressed and contain ^13^C and ^15^N instead of ^12^C and ^14^N. The final protein solution contains about 1 mg/ml protein in 30% v/v acetonitrile (ACN), 0.1% (v/v) trifluoroacetic acid (TFA) in water.

### APT specimen preparation

Supports for the protein sample were made from aluminum wires. These wires were electropolished, first with 20 v/v% nitric acid in methanol at 6 VDC and then with 2 v/v % perchloric acid in 2-butoxyethanol at 1-5 VDC. Consistent results was ensured by a final sharpening and inspection using a dual-beam field emission scanning microscope (SEM) with Xe-plasma focused-ion-beam (PFIB) (FEI Helios, Eindhoven). We did the final sharpening steps in the PFIB using an annular milling pattern with a inner diameter of 150 nm using a beam current of 90 pA at a 16 kV ion accelerating voltage. All specimens had a diameter between 40-100 nm. Finally, an APT measurement of 1-2 million ions was performed (LEAP 5000 XS, laser pulsing at 125 kHz and with 15 pJ pulse energy, 50 K specimen temperature, detection rate 0.2% ions/pulse) to prepare a smooth surface, free from Xe-contamination. This ensured that each Al-specimen would run again when coated with protein.

### Deposition of protein on Al-specimen

A*β*1-42 was deposited on each Al-specimen by slowly dipping in the protein solution. Initially, it was planned to deposit the protein by di-electrophoresis^62^ however we observed that deposition also took place without applying a specimen bias. After deposition, we dried our sample specimen in a desiccator for at least 36h to avoid outgassing in the ultrahigh vacuum chamber of the APT microscope. Different deposition times of 2, 5 and 10 min were used, but there was no difference visible in APT measurements.

**Table I.** Possible overlaps in ion identity in mass-spectra-gained gained from protein deposited on Al-specimen. Peaks in the mass-to-charge-spectra gained from APT measurements. Possible overlaps at different positions in the spectra between different types of ions from the protein and the buffer solution from the protein (Solvent).

| *Da* | Solvent/Chamber ($^{12}\text{C}/^{14}\text{N}$) | | | Protein ($^{13}\text{C}/^{15}\text{N}$) | | | |
| --- | --- | --- | --- | --- | --- | --- | --- |
| 1 | H |  |  | H |  |  |  |
| 2 | H |  |  | H |  |  |  |
| 3 | H |  |  | H |  |  |  |
| 9 | Al |  |  |  |  |  |  |
| 12 |  |  |  | C |  |  |  |
| 13 |  | CH |  | C |  |  |  |
| 13.5 | Al |  |  |  |  |  |  |
| 14 |  | CH<sub>2</sub> | N | CH |  |  |  |
| 15 |  | CH<sub>3</sub> |  | CH<sub>2</sub> | N |  |  |
| 16 |  |  |  | CH<sub>3</sub> | NH | O |  |
| 17 |  |  |  |  | NH<sub>2</sub> | OH |  |
| 18 |  |  |  |  | NH<sub>3</sub> | H<sub>2</sub>O |  |
| 19 |  |  |  |  |  | H<sub>3</sub>O |  |
| 26 |  | C<sub>2</sub>H<sub>2</sub> | CN | C<sub>2</sub> |  |  |  |
| 27 | Al | C<sub>2</sub>H<sub>3</sub> |  | C<sub>2</sub>H |  |  |  |
| 28 | AlH |  | CO | C<sub>2</sub>H<sub>2</sub> | CN |  |  |
| 29 |  |  |  | C<sub>2</sub>H<sub>3</sub> | CNH | CO |  |
| 30 |  |  |  | C<sub>2</sub>H<sub>4</sub> | CNH<sub>2</sub> | COH | CSH |
| 31 |  |  |  | C<sub>2</sub>H<sub>5</sub> | CNH<sub>3</sub> | COH<sub>2</sub> | CSH<sub>2</sub> |
| 32 |  |  |  | O<sub>2</sub> | CNH<sub>4</sub> | COH<sub>3</sub> | S<sub>2</sub> |
| 33 |  |  |  |  | CNH<sub>5</sub> |  |  |
| 40 |  |  |  | C<sub>3</sub>H |  |  |  |
| 41 |  | C<sub>2</sub>NH<sub>3</sub> |  | C<sub>3</sub>H<sub>2</sub> | C<sub>2</sub>N |  |  |
| 42 |  |  |  | C<sub>3</sub>H<sub>3</sub> | C<sub>2</sub>NH | C<sub>2</sub>O |  |
| 43 | AlO |  |  | C<sub>3</sub>H<sub>4</sub> |  | C<sub>2</sub>HO |  |
| 44 | AlOH |  | CO<sub>2</sub> | C<sub>3</sub>H<sub>5</sub> | CNO |  |  |
| 45 |  |  | CO<sub>2</sub>H | C<sub>3</sub>H<sub>6</sub> | CNHO | CO<sub>2</sub> |  |
| 46 |  |  |  | C<sub>3</sub>H<sub>7</sub> | CNOH<sub>2</sub> | CO<sub>2</sub>H |  |
| 47 |  |  |  |  |  | CO<sub>2</sub>H<sub>2</sub> |  |
| 53 |  |  |  | C<sub>4</sub>H |  |  |  |
| 54 |  |  |  | C<sub>4</sub>H<sub>2</sub> |  |  |  |
| 55 |  |  |  | C<sub>4</sub>H<sub>3</sub> |  |  |  |
| 56 |  |  |  | C<sub>4</sub>H<sub>4</sub> |  |  |  |
| 57 |  |  |  | C<sub>4</sub>H<sub>5</sub> |  |  |  |
| 58 |  |  |  | C<sub>4</sub>H<sub>6</sub> |  |  |  |
| 59 |  |  |  | C<sub>4</sub>H<sub>7</sub> | C<sub>2</sub>NOH |  |  |
| 60 |  |  |  | C<sub>4</sub>H<sub>8</sub> |  |  |  |
| 61 |  |  |  | C<sub>4</sub>H<sub>9</sub> |  |  |  |

### Atom probe tomography

APT measurements were performed on a LEAP 5000 XS microscope (Cameca Instruments, Madison, WI) in laser-pulsing mode.The instrument has a ultraviolet laser (UV-laser). The detector efficiency is 81%. We used a detection rate of 2 ions per 1000 pulses and specimens are maintained at 50 K in all experiments. Variable parameter values for pulse energy and pulse rate were used, clarified in the results section.

### Reconstruction

**Figure 7.**
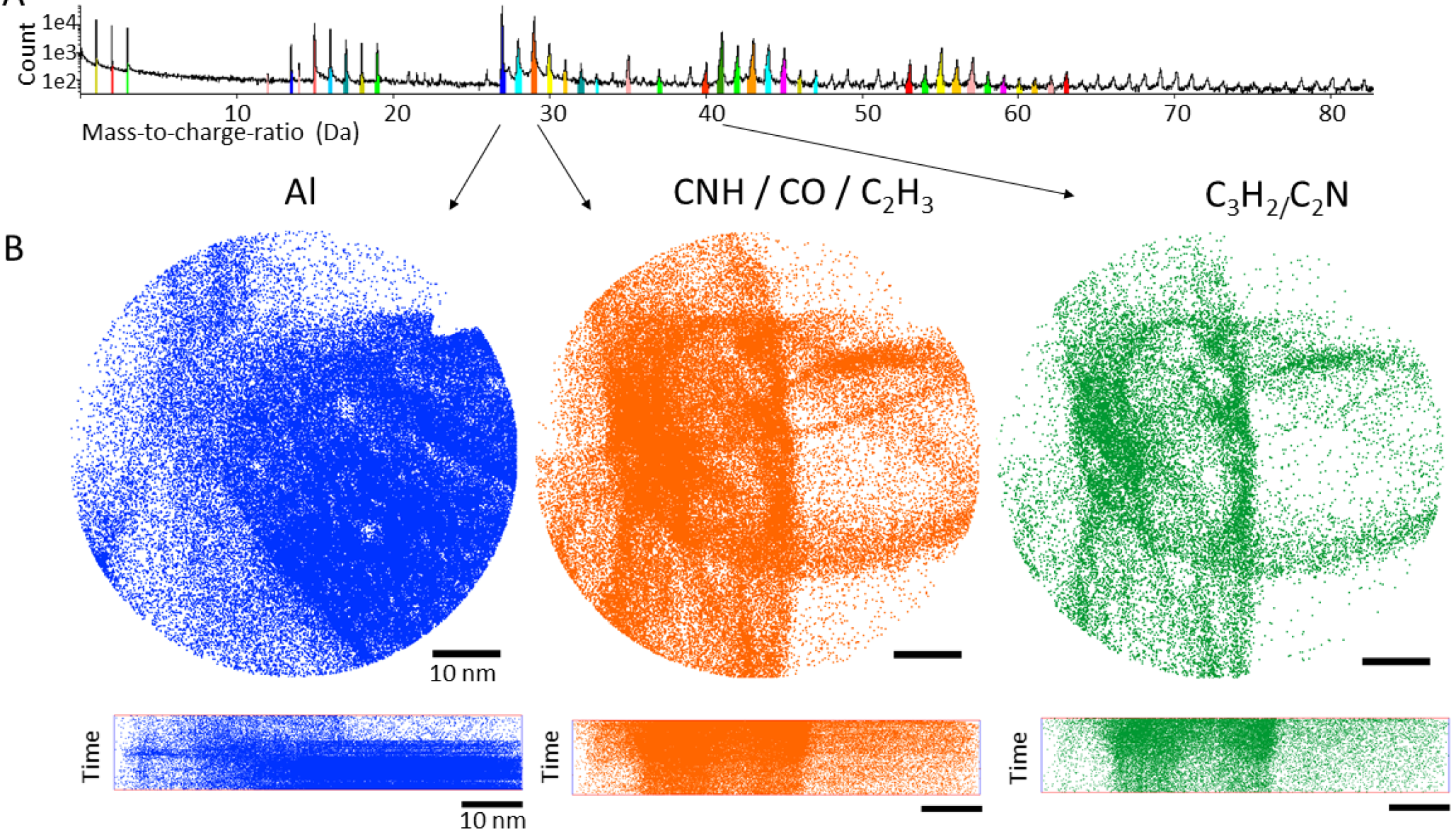
Mass spectrum of an APT measurement and assignment to ion types. A: Mass-to-charge Spectrum for the experiment, showing different ion species in the sample in different colors. B: Distribution of the individual ion species on the detector, top view and side view. From left to right: blue: Peak at 27 Da, Aluminium or C_2_H; orange: Peak at 29 Da could be CNH, CO or C_2_H_3_; green: Peak at 41 Da might be C_3_H_2_ or C_2_N.

APT is primarily a mass-spectrometry technique that also has 3D-reconstruction capabilities. For the first step of reconstruction, IVAS 3.6.14 (Cameca Instruments, Madison, WI) was used. For time-of-flight calibrations, the mass-to-charge peaks at 1.00 Da (H), 13.5 Da (Al), 15.00 (N), 16.00 (O) and 27.00 (Al) were used. For all reconstructions, IVAS default parameters were used. The spatial reconstruction of a field-of-view in X, Y and Z is possible through successive field evaporation of the surface ions^63^. In case of protein deposition, we are especially interested in a reconstruction of the organic surface areas. Organic materials are a challenge for reconstruction algorithms because of differences in the evaporation field in complex materials^64–66^. Therefore we used an approach which takes Z as a sequence number of events on the detector map, with no scaling applied to the depth. A so-called “image stack reconstruction” was formed using the extended position (EPOS) file exported from IVAS using only the singly-detected ions, these being the ions that could be confidently ranged.

## Acknowledgements

Our project to develop new applications for atom probe tomography was founded by the Volkswagen Stiftung through the “Experiment” scheme. We also want to thank Uwe Tezins and Andreas Sturm for help with all issues concerning the measurement instruments.

## Author contributions statement

KAK.R., D.R., D.W., and B.G. conceived the experiment(s), KAK.R., LT.S., A.S. and L.G. conducted the experiment(s), KAK.R., LT.S. and B.G. analysed the results. All authors discussed and reviewed the manuscript.

## Additional information

**Competing financial interests** The authors declare no competing financial interests.

